# A difficult coexistence: resolving the iron-induced nitrification delay in groundwater filters

**DOI:** 10.1101/2024.02.19.581000

**Authors:** Francesc Corbera-Rubio, Emiel Kruisdijk, Sofia Malheiro, Manon Leblond, Liselotte Verschoor, Mark C.M. van Loosdrecht, Michele Laureni, Doris van Halem

**Author notes:** Corresponding author*; Delft University of Technology, Stevinweg 1, 2628 CN Delft, the Netherlands. Notes* The authors declare no competing financial interest.

## Abstract

Rapid sand filters (RSF) are an established and widely applied technology for the removal of dissolved iron (Fe^2+^) and ammonium (NH_4_ ^+^) in groundwater treatment. Most often, biological NH ^+^ oxidation is delayed and starts only upon complete Fe^2+^ depletion. However, the mechanism(s) responsible for the inhibition of NH_4_ ^+^ oxidation by Fe^2+^ or its oxidation (by)products remains elusive, hindering further process control and optimization. We used batch assays, lab-scale columns, and full-scale filter characterizations to resolve the individual impact of the main Fe^2+^ oxidizing mechanisms and the resulting products on biological NH_4_ ^+^ oxidation. Modelling of the obtained datasets allowed to quantitatively assess the hydraulic implications of Fe^2+^ oxidation. Dissolved Fe^2+^ and the reactive oxygen species formed as byproducts during Fe^2+^ oxidation had no direct effect on nitrification. The Fe^3+^ oxides on the sand grain coating, commonly assumed to be the main cause for inhibited nitrification, seemed instead to enhance nitrification by providing additional surface area for biofilm growth. Modelling allowed to exclude mass transfer limitations induced by accumulation of iron flocs and consequent filter clogging as the cause for delayed nitrification. We unequivocally identify the inhibition of NH ^+^oxidizing organisms by the Fe^3+^ flocs generated during Fe^2+^ oxidation as the main cause for the commonly observed nitrification delay. The addition of Fe^3+^ flocs inhibited NH ^+^ oxidation both in batch and column tests, and the removal of Fe^3+^ flocs by backwashing completely re-established the NH_4_ ^+^ removal capacity, suggesting that the inhibition is reversible. In conclusion, our findings not only identify the iron form that causes the inhibition, albeit the biological mechanism remains to be identified, but also highlight the ecological importance of iron cycling in nitrifying environments.

**Graphical abstract:** 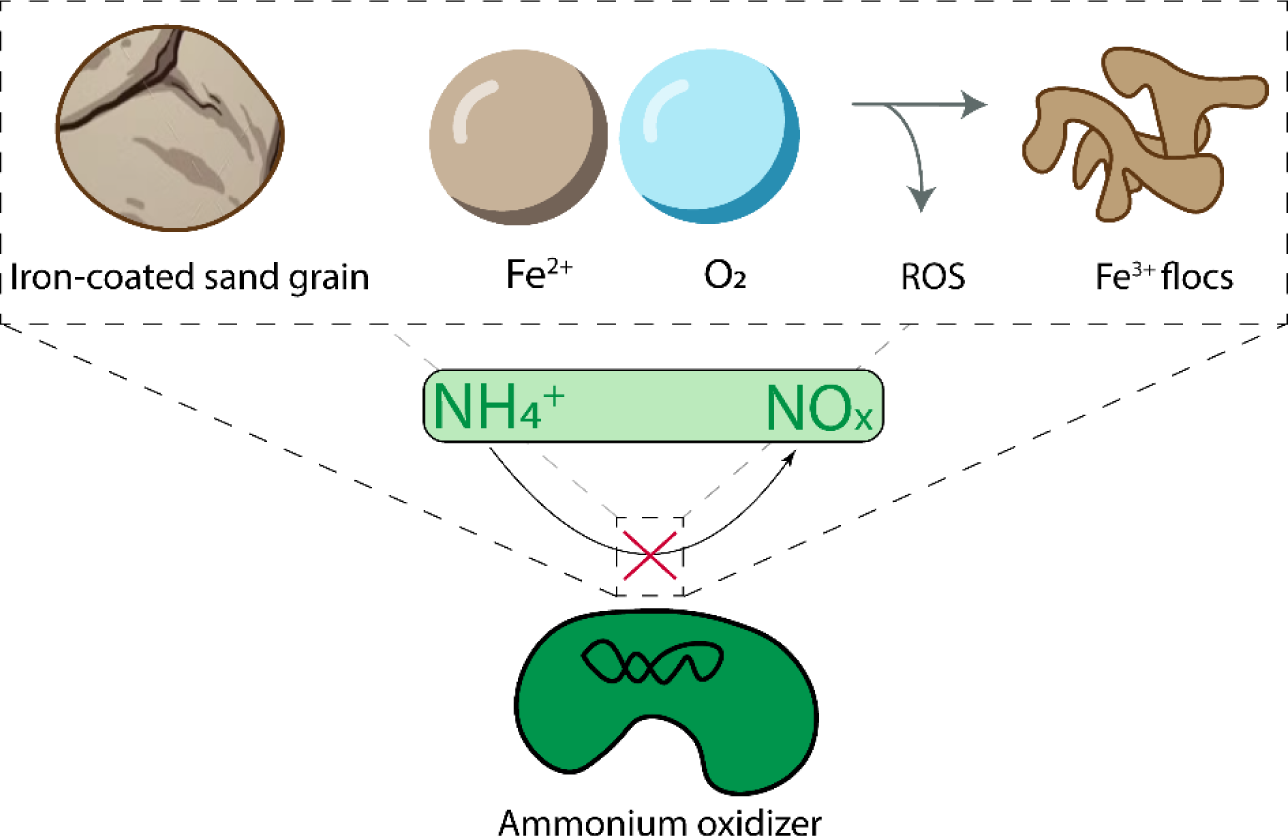

**Highlight:**

- Dissolved Fe^2+^ and reactive oxygen species do not affect NH ^+^ oxidation
- Fe oxide coating aids sand grain colonization by NH ^+^-oxidizing bacteria
- Fe^3+^ flocs inhibit NH ^+^ oxidation by reducing the nitrifying capacity of AOB
- Changes in transport patterns due to clogging do not play a major role in NH ^+^ oxidation
- The inhibition of NH_4_^+^ oxidation is reversible and reduced by backwashing

## 1. Introduction

Anaerobic groundwater is an excellent drinking water source due to its microbiological safety, stable temperature, and composition (1–3). Rapid sand filters (RSFs), preceded by an aeration step, are the most commonly applied technologies for the removal of major groundwater contaminants such as dissolved iron (Fe^2+^) and ammonium (NH_4_ ^+^) (4). Complete segregation of the removal processes is commonly observed in full-scale systems, where Fe^2+^ is removed at the filter top and NH_4_ ^+^ oxidation starts only upon Fe^2+^ depletion (5–7). Despite decades of research and practice, the mechanisms governing the stratification of Fe^2+^ and NH_4_ ^+^ removal remain elusive, leading to poor process control, filter over-dimensioning, and even incomplete ammonium removal (8). A clear understanding of this phenomenon is key to improve current RSF operation and for the design of novel resource-efficient systems. Fe^2+^ removal in RSFs is a two-step process that consists of (i) Fe^2+^ oxidation to Fe^3+^ and (ii) subsequent entrapment of the oxidation products within the filter bed. Fe^2+^ oxidation can take place via three different oxidation mechanisms: homogeneous (flocculent), heterogeneous (surface-catalytic), and biological (9). Homogeneous oxidation is a chemical reaction between dissolved Fe^2+^ and dissolved oxygen. Poorly crystalline, low-density hydrous ferric oxide flocs (Fe^3+^ flocs) with large specific surface area are formed (10,11), with reactive oxygen species (ROS) derived from Fenton chemistry as byproducts (12). Fe^3+^ flocs accumulate within the pore space of RSFs during operation, clogging the filter and thus forcing backwashing. Heterogeneous oxidation occurs when Fe^2+^ adsorbed onto the surface of sand grains is oxidized. Fe^2+^-surface complex formation is followed by the oxidation of Fe^2+^ to Fe^3+^ and subsequent hydrolysis, producing compact hydrous ferric oxides that generate a coating on the sand grains, contributing to sand grain growth (11). Biological iron oxidation is catalyzed by iron-oxidizing bacteria (FeOB). FeOB and its Fe oxidation products form a porous coating on the sand grains, which may release stalk-like ferric oxides into the pore water. On the contrary, NH_4_ ^+^ oxidation is an exclusively biological process known as nitrification and carried out by NH_4_ ^+^-oxidizing (AOB) and comammox bacteria, as well as NH_4_ ^+^-oxidizing archaea, with nitrite and nitrate as products (13). During the last decades, virtually all compounds that play a role in Fe^2+^ oxidation within RSFs have been reported to inhibit NH_4_ ^+^ oxidation.

Dissolved Fe^2+^ and ROS have been shown to decrease the growth rate of pure cultures (14,15), and similar conclusions were drawn for Fe^3+^ flocs (16) and Fe oxide coating (17,18), the products of homogeneous and heterogeneous iron oxidation respectively. In a real-life industrial set-up, de Vet and colleagues (2009) concluded that the presence of Fe^3+^ flocs and Fe oxide coating reduces the nitrification capacity of full-scale sand filters. While this extensive body of knowledge certainly contributes to our understanding of the interactions that take place, it also poses a challenge in distilling the essential information - *which are the specific processes responsible for the segregation between Fe^2+^ and NH_4_ ^+^ oxidation in RSFs.* Here, we aim at determining which compound(s) involved in Fe^2+^ oxidation have a major impact on NH_4_ ^+^ oxidation in RSFs. A systematic and quantitative evaluation of the impact of each compound on nitrification has been adopted, including dissolved Fe^2+^, Fe^3+^ flocs, ROS, and Fe oxide coating. Each Fe^2+^ oxidizing mechanism and the involved compound was separately studied using a combination of batch essays, laboratory columns, and full-scale RSFs observations, supported by modelling.

## 2. Materials and Methods

### 2.1. Full-scale filter sampling and media collection

Fe oxide-coated filter media was collected from the top of a filter located in DWTP Hammerflier, operated by Vitens, in Den Ham (the Netherlands). The filter is fed with aerated groundwater with 2.6mg NH_4_-N^+^·L^-1^ and 3 mgFe^2+^·L^-1^. NH_4_ ^+^ removal is completely absent at the filter top, and only starts after Fe is removed (Figure S1). Filter media was transported in water at 4 °C and added to the lab-scale column within 8h.

Full-scale sand filter experiments (Results 3.6) were performed in a groundwater-fed rapid sand filter located in DWTP Loosdrecht, operated by Vitens, in Loosdrecht (the Netherlands). The filter consisted of a 1 m top layer of anthracite (1.4-2.0 mm), and a 1 m bottom layer of sand (0.8-1.25 mm), which were slightly mixed. Water samples were taken before and after backwashing. The first round of water samples were obtained after a filter run time of about 52 hours with a flow of about 225 m^3^/hour. Afterwards, the filter was backwashed following the drinking water companies standard protocol. A second round of water samples was taken after about 2.5 hours operation.

In both cases, the filters were equipped with water taps which enabled water sampling along the filter depth of the filter bed as well as in the supernatant water. Water samples were immediately filtered (0.45 µm), stored at 4 °C, and measured within 24 h as explained below.

### 2.2. Laboratory column experiments

Six sand filter columns of 20 cm height and an inner diameter of 2.5 cm were operated continuously in down-flow mode at a superficial velocity of 1.5 m·h^-1^ with growth medium (L^-1^; 2 mg NH_4_-N, 17.4 mg K_2_HPO_4_, 60µL trace elements solution (L^-1^; 15 g EDTA, 4.5 g ZnSO_4_ ·7H_2_O, 4.5 g CaCl_2_·2H_2_O, 3 g FeSO_4_·7H_2_O, 1 g H_3_BO_3_, 0.84 g MnCl_2_·2H_2_O, 0.3 g CoCl_2_·6H_2_O, 0.3 g CuSO_4_·5H_2_O, 0.4 g Na_2_MoO_4_·2H_2_O, 0.1 g KI)) (Soler-Jofra et al., 2018) at pH 7.5 ± 0.1 and ambient temperature (20°C). pH and dissolved oxygen were continuously sensed (Applikon AppliSens, the Netherlands) at the influent and effluent of the column. The filter medium consisted of either (*i*) 240 g of virgin quartz sand (Aqua-Techniek B.V., the Netherlands), (*ii*) 190 g of Fe oxide-coated sand from the top of a full-scale filter in DWTP Hammerflier, or (*iii*) a mixture of 120 g virgin sand and 100 g Fe oxide-coated sand to achieve a bed height of 10 cm (Figure 1).

**Figure 1.**
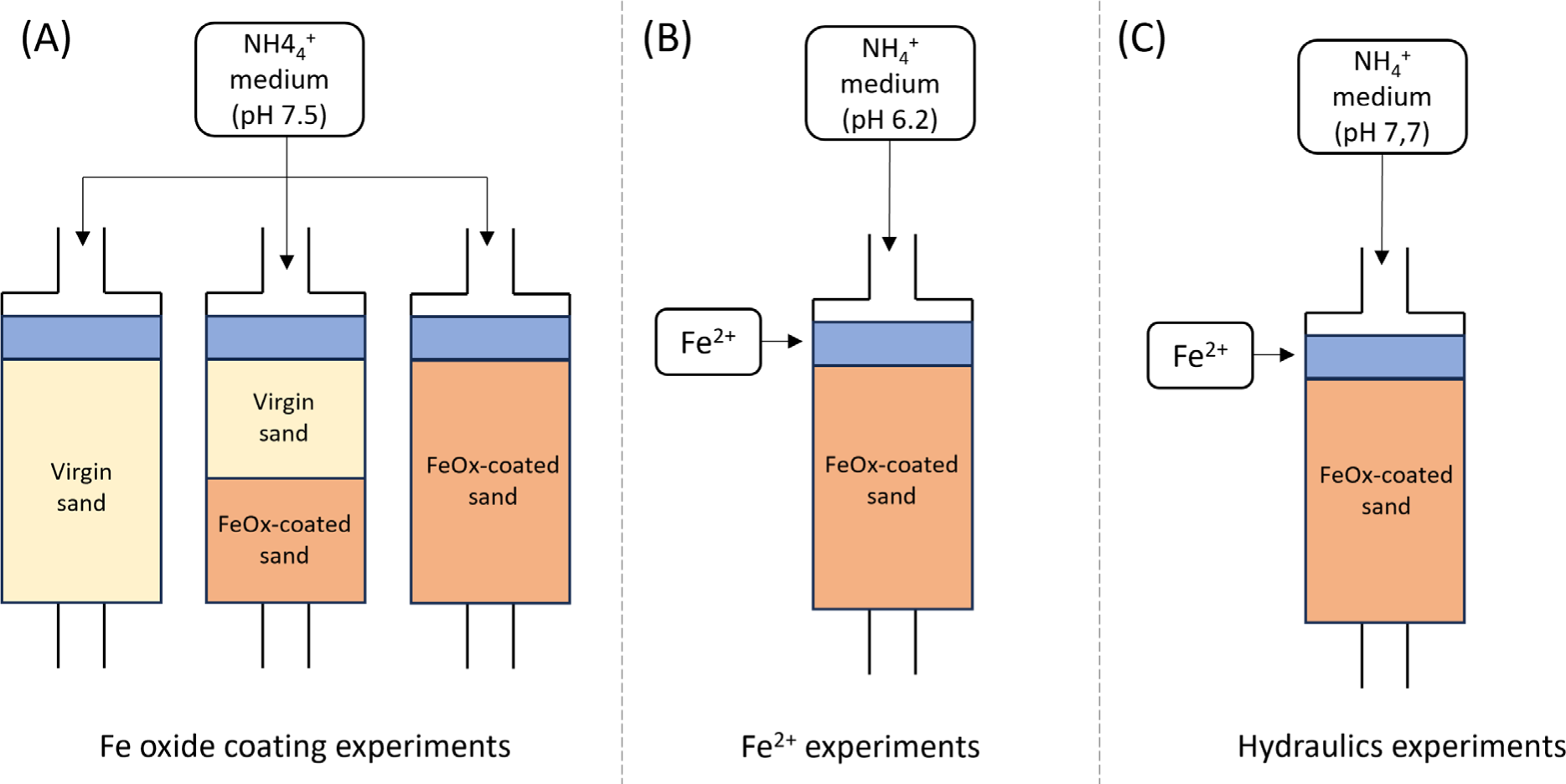
Schematic representation of the column experiments. A) Fe oxide coating experiments. B) Fe^2+^ experiments. C) Hydraulics experiments.

#### 2.1.1. Fe oxide coating experiments

For the Fe oxide coating experiments (Results 3.1), filter media from a full-scale filter with an NH_4_ ^+^-oxidizing biofilm was used to inoculate the columns as previously reported (20). In short, 0.5 g of sand was vortexed with 10 mL tap water for 1.5 minutes. A total of 100 g of sand were processed. The resulting solution was amended with ammonium to reach a concentration of 10 mgN-NH_4_ ^+^·L^-1^ and was recirculated through each column in a closed-circuit mode for 25 days. 250 mL of nitrifying sludge (4.6g VSS·L^-1^) were added to the system on days 10^th^ (and 20^th^ for the virgin sand column), as done in previous research (21). Before switching to standard operation (*i.e.* not recirculating), all columns were backwashed with 13.6L·h^-1^ tap water for 9 min to remove suspended particles.

#### 2.1.2. Fe^2+^ and hydraulics experiments

For the Fe^2+^ experiments (Results 3.3), the microbial community of a column filled with 190 g of Fe oxide-coated sand from the top of a full-scale filter in DWTP Hammerflier was adapted to pH 6.2 by step-wise decreasing the influent pH throughout a period of 2 months. For the hydraulics experiments (Results 3.4), a column filled with 190g of Fe oxide-coated sand from the top of a full-scale filter in DWTP Hammerflier was operated at pH 7.7 throughout a period of 2 months to reach complete, stable NH_4_ ^+^ removal. In both cases, an Fe^2+^ solution (0.5 g FeCl_2_·4H_2_O·L^-1^) was fed from one of the lateral valves, directly into the supernatant, to reach a concentration of 1.5 mgFe^2+^·L^-1^.

### 2.3. Batch experiments

The maximum NH ^+^ removal rates of the filter media were determined in batch. 4 g of wet filter media, 50 mL of tap water, 100 μL of trace element solution (L^-1^; 15 g EDTA, 4.5 g ZnSO_4_ ·7H_2_O, 4.5 g CaCl_2_·2H_2_O, 3 g FeSO_4_·7H_2_O, 1 g H_3_BO_3_, 0.84 g MnCl_2_·2H_2_O, 0.3 g CoCl_2_·6H_2_O, 0.3 g CuSO_4_·5H_2_O, 0.4 g Na_2_MoO_4_·2H_2_O, 0.1 g KI) and 500 µL of 100 nM K-phosphate buffer solution (1.37 g KH_2_PO_4_ ·L^-1^, 1.75 g K_2_HPO_4_·L^-1^, pH 7.5) were mixed in 300mL shake flasks. After an acclimatization period of 30 min at 25 °C and 100 rpm, each flask was spiked with 1 mL of feed (100 mgN·L^-1^ NH_4_Cl and 87mg K_2_HPO_4_ ·L^-1^) (Sigma Aldrich, Saint Louis, Missouri USA). Liquid samples (1 mL) were taken at different time intervals throughout the entire process. Maximum removal rates per mass (wet weight) of filter media were calculated by fitting the concentration profiles to a first-order kinetic rate equation. To study the effect of ROS, the batch tests were amended with 75 µL of ultrapure water (control), 4 g Fe^2+^·L^-1^ (Fe^2+^) or 4 g Fe^3+^-flocs·L^-1^ (Fe^3+^) and 254 µL of 4 g·L^-1^ TEMPOL, 215 g·L^-1^ mannitol, 0.4 g·L^-1^ catalase (3000 U·mg^-1^), 32 g·L^-1^ methanol or ultrapure water every 30 min during 3 h, mimicking the continuous operation of full-scale rapid sand filters. TEMPOL was added at a 1:1 TEMPOL:Fe^2+^ molar ratio (22), methanol at 200:1, in the range tested by Tong et al. (2022), and mannitol and catalase to reach a final concentration of 360 U·mL^-1^ and 30 mM, similar to the approach of Brinkman et al. (2016). Similarly, to test the effect of Fe^3+^ flocs 75 µL of 4/8/12/16 g Fe^3+^-flocs·L^-1^ (Fe^3+^) were added every 30 min during 2.5h.

The Fe^2+^ solution was prepared anaerobically in a vinyl PVC anaerobic chamber (Coy Laboratory Products, Grass Lake, Michigan, USA) at pH 2 to avoid Fe^2+^ oxidation. To prepare the Fe^3+^ floc solution, a concentrated solution of Fe^2+^ was slowly added into a recipient vessel with ultrapure water vigorously agitated to ensure direct Fe^3+^ floc formation.

### 2.4. Analytical procedures and data analysis

Samples for ammonium, nitrite, and nitrate quantification were immediately filtered through a 0.2 µm nanopore filter and measured within 12 h using photometric analysis (Gallery Discrete Analyzer, Thermo Fischer Scientific, Waltham, Massachusetts, USA). Samples for iron were analysed within 12 h by spectrophotometry (DR3900, Hach Company, Ames, Iowa, USA) using the LCK320 kit (Hach Company, Ames, Iowa, USA). Mn was analysed using inductively coupled plasma mass spectrometry (ICP-MS, Analytikal Jena PlasmaQuant MS). Water pressure along the column height was measured with a GMH 3100 manometer (Greisinger, Germany).

Batch test data were plotted and statistically analysed with GraphPad Prism software (Dotmatics, UK), which uses a statistical method equivalent to ANCOVA (Analysis of Covariance) to determine if the slopes between two linear regressions are significantly different. A significant difference was considered when p-value < 0.05.

Hydraulic retention time was measured using salt as a tracer. 10 mL of 1 g/L NaCl were spiked to the supernatant of the column. The increase in electric conductivity in the water effluent was measured using a flow-through cell connected to an electrical conductivity sensor (WTW, Xylem Analytics, Weilheim, Germany).

The residence time of groundwater in the filter was estimated to assess oxidation over time in the filter and to obtain kinetic rate constants of the reactants. The height of the sampling location was converted to a residence time by calculating the volume of water within the supernatant and filter bed based on the height and an assumed porosity of 0.4 and dividing the volume of water by the flow to obtain the residence time. First-order rate constants were estimated by fitting a first-order rate equation to the observed concentrations over residence time in the supernatant and filter bed. This was done using a least-squares routine in Python (v. 3.6.4).

## 3. Results

### 3.1. Fe oxide coating does not inhibit NH_4_ ^+^ oxidation

Fe oxide-coated media contain reactive surfaces that might hamper the colonization, growth, or NH_4_ ^+^ oxidation capacity of NH_4_ ^+^-oxidizing organisms. To test this hypothesis, three lab-scale columns with either (i) virgin sand, (ii) Fe oxide-coated sand or (iii) 50% virgin – 50% Fe oxide-coated sand were inoculated and run in continuous mode for 45 days after a 25-day inoculation in 100% recirculation mode (Figure 2A). The two columns with Fe oxide-coated sand started consuming NH_4_ ^+^ immediately after inoculation and achieved complete removal after 10 (Fe oxide-coated sand) and 26 days (mixed sand). On the contrary, NH_4_ ^+^ conversion was absent in the virgin sand column at the beginning of the experiment, and a second inoculation with nitrifying activated sludge (day 10) did not yield better results. Therefore, on day 20, the effluent of the Fe oxide-coated sand filter was connected to the influent of the virgin sand column. Complete NH_4_ ^+^ removal was achieved within 5 days, presumably due to suspended cells carried from the Fe oxide-coated sand filter. Once the effluent of the Fe oxide-coated sand filter was disconnected (day 25), NH_4_ ^+^ consumption decreased rapidly and stabilized at 12 ± 3%. Microscopic visualization of the media showed that the onset of NH_4_ ^+^ consumption coincided with the appearance of Fe oxides (Figure 2C) on the previously virgin sand (Figure 2B).

**Figure 2.**
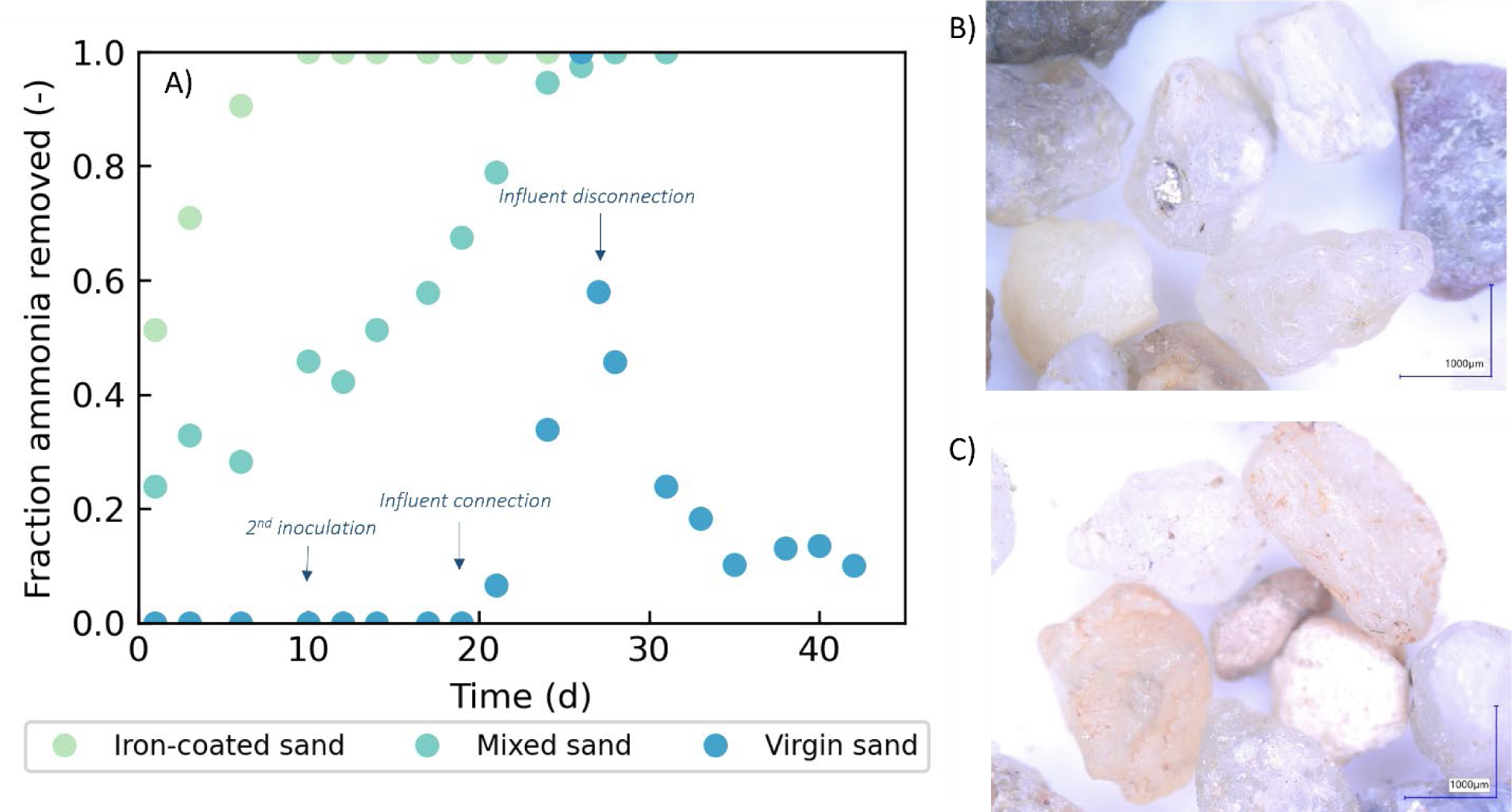
A) Fraction of NH_4_^+^ removed in the effluent of the three lab-scale sand filters during continuous operation. Blue arrows indicate the second inoculation (day 10) and the connection and disconnection of the effluent of the Fe-oxide coated filter to the influent of the virgin sand filter (days 20 and 25). B) Microscopic image of the virgin sand on day 10. C) Microscopic image of the virgin sand on day 35, slightly covered with Fe oxides.

Overall, these results show not only that Fe oxides do not hamper NH_4_ ^+^ removal, but that they play an essential role in sand grain colonization by NH_4_^+^-oxidizing organisms.

### 3.2. Fe^2+^-derived ROS do not inhibit NH_4_^+^ oxidation

Reactive oxygen species (ROS) are formed as by-products of homogeneous Fe oxidation (24). The potential inhibition of NH_4_ ^+^ oxidation by ROS was tested in batch tests with Fe oxide-coated sand with NH_4_ ^+^-oxidizing activity amended with ROS-quenchers/scavengers TEMPOL, mannitol, catalase, and methanol. Negative control experiments showed that ROS quenchers/scavengers do not diminish the rate of NH_4_ ^+^ oxidation (light green). On the contrary, Fe^2+^ addition (which quickly oxidized to Fe^3+^, dark green) significantly decreased the rate, regardless of the addition of ROS-quenchers/scavengers (-24.8 ± 1.8% on average). Adding Fe^3+^ flocs instead of Fe^2+^ resulted in a similar level of inhibition (-30%, blue). Altogether, these results prove that ROS do not hamper NH_4_ ^+^ oxidation on Fe oxide-coated filter sand, and suggest that Fe^3+^ flocs are the main inhibitor of NH_4_ ^+^ oxidation.

### 3.3. Fe^2+^ presence does not interfere with NH_4_ ^+^ oxidation

Dissolved Fe^2+^ is the Fe form that enters full-scale sand filters. To evaluate its effect on NH_4_ ^+^ oxidation, a lab-scale column filled with Fe oxide-coated sand was continuously fed with NH_4_ ^+^-amended tap water at pH 6.2. Complete NH_4_ ^+^ removal was obtained after 2 months, after which Fe^2+^ addition into the supernatant started. The low pH allowed Fe^2+^ to penetrate deep into the bed without oxidizing (Figure 4A), as both homogeneous and heterogeneous oxidation rates substantially decrease with decreasing pH (25,26). In accordance, Fe^3+^ was barely observed throughout the filter (Figure 4B), leaving adsorption as the only cause of Fe^2+^ removal. The NH_4_ ^+^ concentration profile along the filter depth remained relatively stable during the course of the experiment (Figure 4C), demonstrating that the presence of (adsorbed) Fe^2+^ cannot explain the inhibition of NH_4_ ^+^ removal observed at the top section of full-scale sand filters.

### 3.4. Fe^3+^ flocs spatially delay NH_4_^+^ oxidation

Similarly to section 3.3, a lab-scale sand filter filled with Fe oxide-coated sand was continuously fed with tap water amended with NH_4_ ^+^ but now at higher pH (pH=7.7). Full NH_4_ ^+^ removal was obtained after 2 months of acclimation, after which Fe^2+^ addition into the supernatant started. Fe^2+^ was consistently oxidized at the filter top, with >75% removal in the first 5 cm (Figure 4D). Homogeneous Fe oxidation was the main Fe removing mechanism in the system, and most flocs accumulated at the top (Figure 4E). The accumulation of Fe^3+^ flocs throughout the experiment resulted in filter clogging in the top of the column, which changed the hydraulic conditions of the filter (see SI 8.2) and spatially delayed the removal of NH_4_ ^+^ (Figure 4F). At the end of the experiment, NH_4_ ^+^ oxidation was completely absent in the first 5 cm of the filter bed (Figure 5), and the estimated first-order rate constants (k) decreased in the downstream sections (from 0.022 to 0.015 s^-1^). These results show that Fe^3+^ floc accumulation is directly responsible for the loss of nitrifying activity in sand filters, supporting the findings of Figure 3. In addition, they indicate that the loss in nitrifying activity is more prominent in zones where more Fe^3+^ flocs accumulate, with total loss of activity at high floc content. The following sections aim at understanding the inhibiting mechanism.

### 3.5. Increasing pore velocities due to clogging alone cannot explain the spatial delay in NH_4_ ^+^ oxidation

Changing transport patterns in the filter bed due to Fe^3+^ floc accumulation may affect NH_4_ ^+^ oxidation. Since the sand filter was operated at constant flow, Fe^3+^ floc accumulation (*i.e*., increase in filter bed resistance) directly translated into (local) increases in water pore velocity. Consequently, contact times between AOB and NH_4_^+^ are shorter, which could theoretically push NH_4_^+^ oxidation deeper into the filter bed. To evaluate the contribution of increased pore water velocities to the spatial delay of NH_4_ ^+^ oxidation, we estimated the average pore water velocity needed to explain the four observed NH_4_ ^+^ concentration profiles in Figure 5 assuming that Fe^3+^ flocs do not inhibit AOB *i.e.* maintaining a constant *k* of 0.02s^-1^ for the top 5cm of the filter bed. A local average increase in pore water velocity changes the hydraulic retention time (HRT) in the filter, and can be estimated using the equation below:

where *t* is the travel time between the two depths (s), *C_depth= i cm_* is the NH_4_-N concentration (mg·L^-1^) at a depth *I, and k* is the NH_4_ ^+^ oxidation rate constant (s^-1^).

Simulating the results for the top 5 cm of filter bed for each day yields theoretical HRTs of 23, 19, 14, and ∼0 seconds for day 1, 2, 3, and 4, respectively. This means that a 1.2, 1.7, and infinite fold-increase in the water pore velocity are needed to explain the decrease in NH_4_ ^+^ oxidation solely by changes in HRT (Table 1), which is proportional to a loss in pore space of 17%, 41%, and ∼100%, for Days 2, 3, and 4, respectively. These percentages are substantially higher than the maximum percentage of pore space that could be filled up with the theoretical amount of Fe^3+^ flocs formed during the experiment (1%). We deem it, therefore, unlikely that changes in hydraulic conditions in the clogged column alone can explain the spatial delay in NH_4_^+^ oxidation.

**Table 1.**
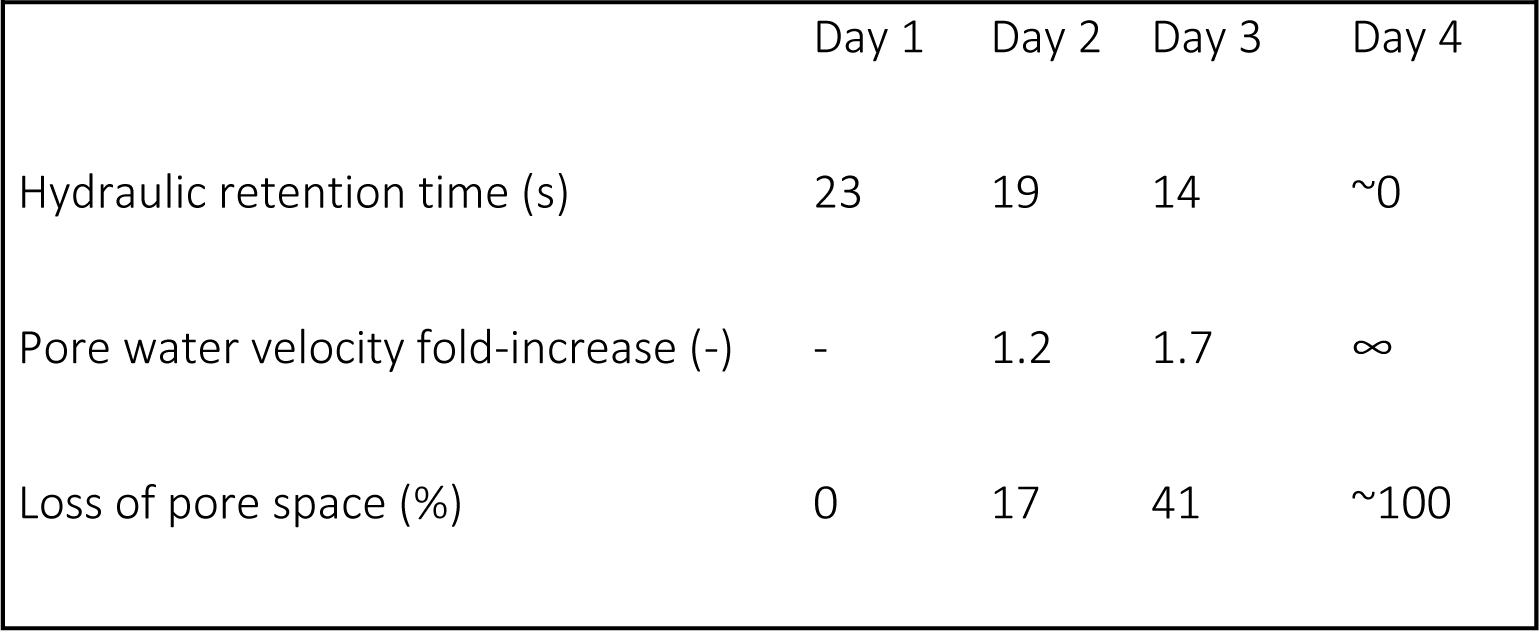
Estimated hydraulic retention time and fold increase in pore water velocity are needed to explain the spatial delay in NH_4_^+^ removal in the lab-scale sand filter solely due to changes in transport phenomena.

### 3.6. Fe^3+^ flocs reduce the activity of NH_4_ ^+^-oxidizing bacteria (AOB)

The potential direct impact of Fe^3+^ flocs on AOB – *i.e.*, reducing their nitrifying capacity - was tested in batch tests. Sand grains from the same NH ^+^ oxidising full-scale sand filter as for the column study were incubated with 0, 30, 60, 90, and 120 mgFe^3+^flocs·L^-1^. Sand grains are continuously mixed and thus clogging and transport limitations do not play a role. Figure 6 shows that higher floc content yielded proportionally lower maximum NH_4_ ^+^ removal rates, demonstrating that Fe^3+^ flocs inhibit AOB.

Comparing the amount of Fe^3+^ flocs accumulated in this experiment and the lab-scale sand filter (Figure 5) allows to estimate the likelihood of Fe^3+^ flocs contributing to the spatial delay in NH_4_^+^ in the sand filter. About 100 mg of Fe^3+^ flocs were produced during the sand filter experiment, which corresponds to a floc content of >2000 mg Fe/L assuming a homogeneous distribution across all pores in the column and total floc retention. A more realistic assumption is that all flocs were entrapped within the first 5 cm of the filter bed (Figure 5e), which yields a Fe^3+^ floc content of >15,000 mg Fe/L. In both cases, the floc content is 1 to 2 orders of magnitude higher than the maximum content of flocs in the batch experiments (∼120 mg/L) (Figure 6), which was enough to reduce the NH_4_ ^+^ removal capacity of the sand grains by 50%. Therefore, we conclude that AOB inhibition by Fe^3+^ flocs is likely the main mechanism inhibiting NH_4_^+^ removal at the top of sand filters.

### 3.7. Reversible inhibition of NH_4_ ^+^ and Mn^2+^ oxidation by Fe^3+^ flocs

The reversibility of the spatial delay in NH_4_ ^+^ removal caused by Fe^3+^ flocs was evaluated by assessing the NH_4_ ^+^ concentration profiles along the filter depth of a full-scale filter before and after backwashing (Figure 7A). NH_4_ ^+^ removal was almost absent at the filter top at the end of a filter run, *i.e.* before backwashing. Backwashing moved the NH_4_ ^+^ removal to the filter top, even though dissolved – Fe^2+^ - and particulate – Fe^3+^ flocs - Fe were not completely removed (see Figure S3).

The effect of backwashing was also evaluated for Mn^2+^, to explore putative parallelisms between the inhibiting effects of Fe^3+^ flocs (Figure 7B and 7C). Similarly, while Mn^2+^ was barely removed at the filter top before backwashing, removing the Fe^3+^ flocs moved its removal to the filter top, resulting in a similar 2.68 and 2.08 fold-increase in the first-order oxidation rate constants for NH_4_ ^+^, and Mn^2+^, respectively. Note that, contrary to NH_4_ ^+^, the removal of Mn^2+^ does not exclusively rely on microorganisms, and can be chemically catalyzed. Interestingly, the trends are nonetheless similar, which may indicate that Fe^3+^ flocs affect the removal of both contaminants in a similar, yet undescribed fashion.

## 4. Discussion

### 4.1. Dissolved Fe^2+^, reactive oxygen species, and Fe oxide coating do not decrease the NH ^+^ removal capacity of sand filters

We evaluated the influence of dissolved Fe^2+^, reactive oxygen species, and Fe oxide coating on NH ^+^ oxidation in sand filters and found no major inhibitory effects (Figures 2, 3, and 4C). Bacteria exposure to Fe^2+^ and ROS have often been associated with detrimental physiological effects. Swanner et al., (2015) showed that Fe^2+^ addition to a cyanobacteria culture decreased growth rate and attributed it to the intracellular accumulation of ROS. In another example, Zhang et al., (2020) found that exposing *Shewanella oneidensis* strain MR-1 to Fe^2+^ minerals and oxygen decreased cell viability and associated it with ROS production as well. Fe^2+^ concentrations, a proxy for ROS concentration under aerobic conditions, were much higher in our batch tests than in those studies (70 mM vs 5 mM and 80 µM, respectively). However, our results clearly show no immediate decrease in the NH_4_ ^+^ removal rate, and effects on cell viability are likely marginal because NH_4_ ^+^ removal capacity in full-scale sand filters recovers immediately after backwashing (Figure 7), proving inhibition to be reversible. We hypothesize that ROS exert selective pressure on the sand filter ecosystem, and as a result, only organisms able to tolerate ROS thrive. An example of adaptation is the increase in catalase activity and upregulation of several antioxidant genes in AOB *Nitrosomonas oligotropha PLL12* upon exposure to ROS observed by Wang et al., (2023).

**Figure 3.**
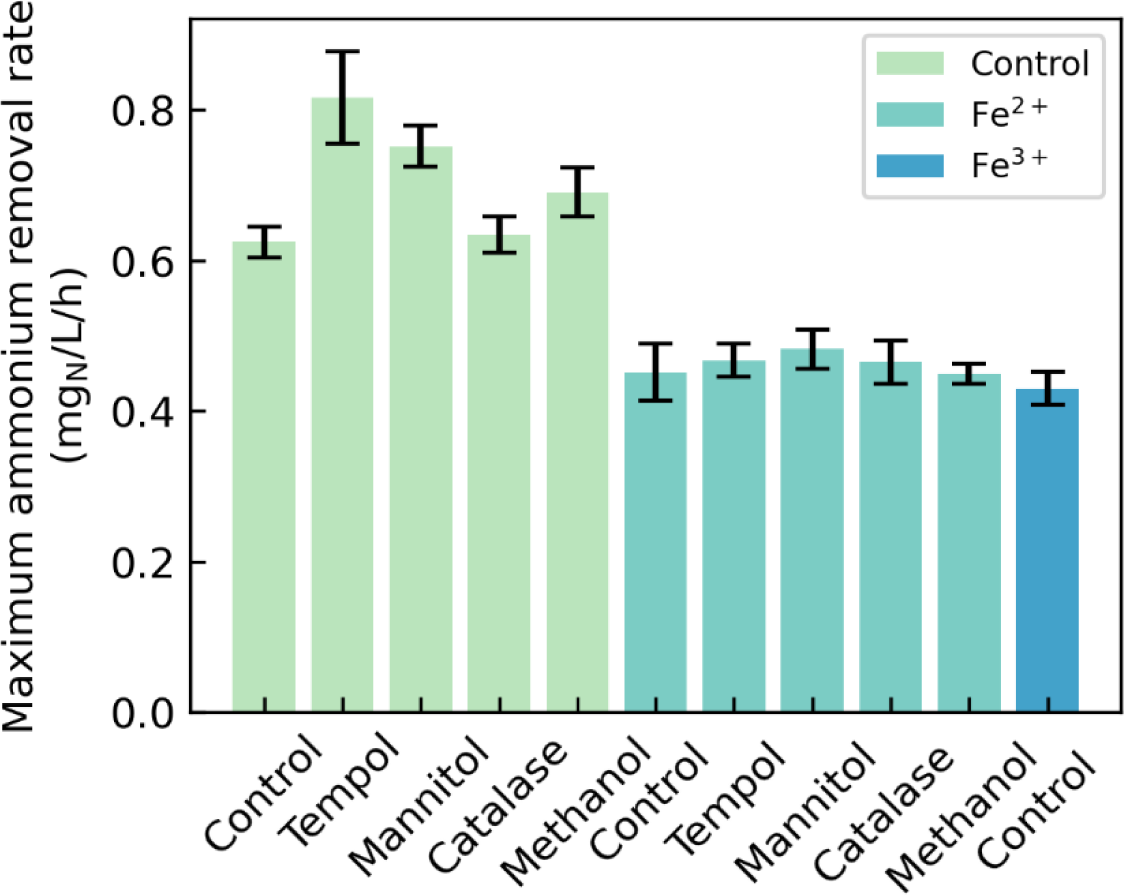
Maximum NH ^+^ removal rates of the sand grains in the absence (light green), or presence of Fe^2+^ (dark green) or Fe^3+^ flocs (blue). Control tests contain no ROS quenchers/scavengers (TEMPOL, mannitol, catalase, and methanol). All tests were performed aerobically to ensure quasi-instantaneous Fe^2+^ oxidation and concomitant ROS formation, mimicking the conditions of full-scale sand filter supernatants. Activities were quantified in triplicates, error bars represent standard deviation.

**Figure 4.**
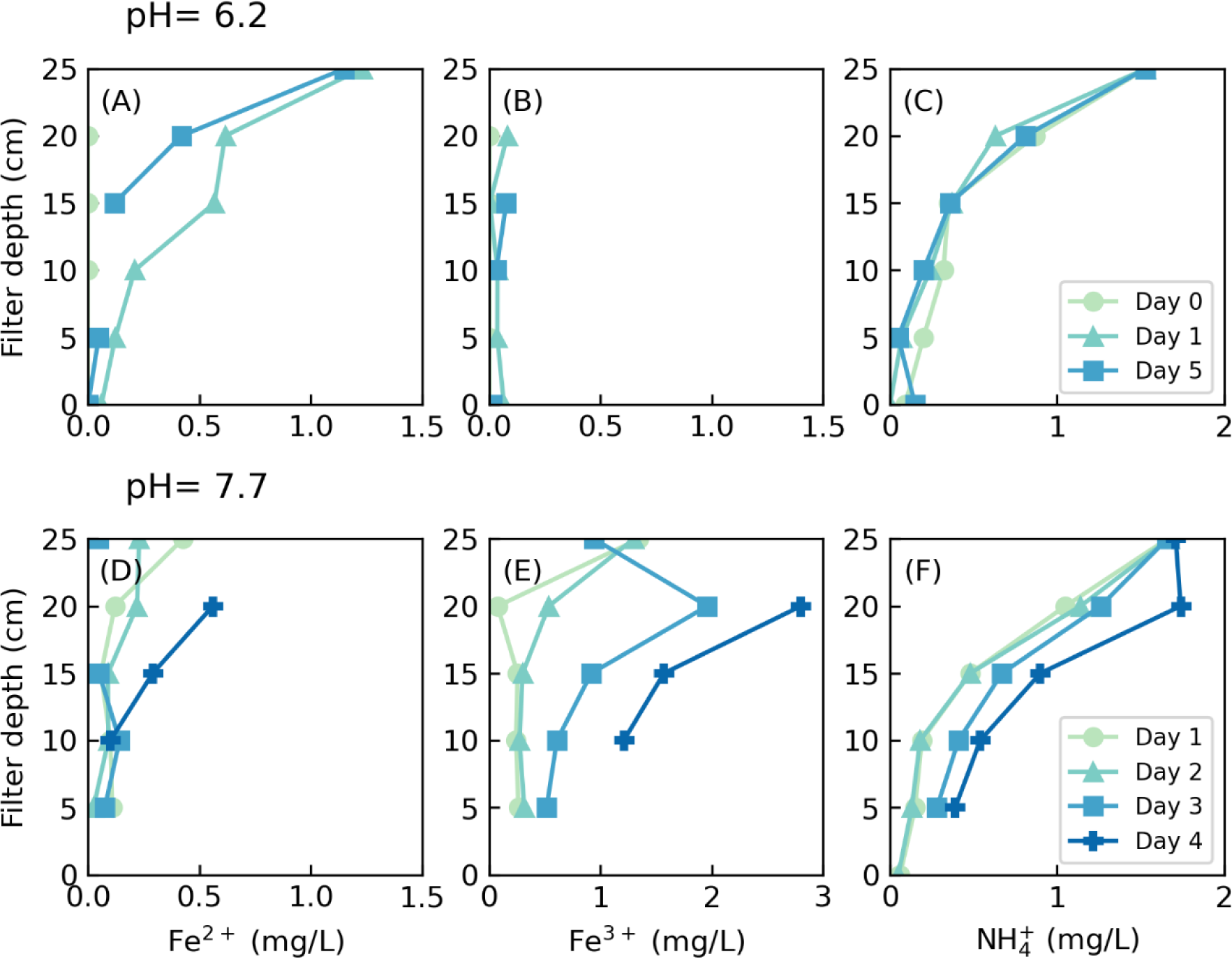
Fe^2+^ (A, D), Fe^3+^ (B, E), and NH_4_^+^(C, F) removal profiles along the depth of a lab-scale sand filter during continuous dosing of Fe^2+^ at pH 6.2 (A, B, C) and 7.7 (D, E, F), respectively. The initial Fe^2+^ concentration is 1.5 mg/L. Data points represent single measurements. Dashed horizontal line indicates supernatant position.

In a similar fashion, several studies on both pure and mixed cultures pointed out the toxic effects of Fe oxides. Growth inhibition by several Fe oxides – including ferrihydrite, the most common Fe oxide in sand filters (28) – was observed by Du et al., (2019), and decreases in cell viability upon exposure to Fe oxide nanoparticles (29) and Fe oxide-containing clay minerals (18) are also commonly reported. Our colonization experiments (Figure 2) not only indicate that Fe oxides do not impede the growth of AOB on sand grains, but also that their presence even accelerates the colonisation process. In addition, Fe oxide coatings have high specific surface areas (30), which seem to be a critical factor for biofilm growth (31). One of the limitations of our study is that the experiments were carried out on a stable Fe oxide layer, *i.e.* no Fe^2+^ was fed into the system, so no new Fe oxides were generated. In contrast, the Fe oxide coating of sand grains of full-scale sand filters continuously grows during operation as a result of heterogeneous Fe^2+^ oxidation. One can argue that this dynamic situation may force AOB to match the growth rate of Fe oxides to ensure nutrient availability and avoid cell encrustation. However, Gülay et al., (2014) observed a positive correlation between cell abundance and mass mineral coating in full-scale sand filter grains, suggesting that the Fe oxide coating enhances, rather than hinders, the NH_4_ ^+^ removal capacity of full-scale sand filters. Overall, we show that dissolved Fe^2+^, reactive oxygen species, and Fe oxide coating do not have a major detrimental effect on AOB activity, contrary to what is commonly reported in literature. Beyond groundwater filtration, these results showcase the limitations of extrapolating results from pure culture studies into complex microbial communities.

### 4.2. Direct inhibition of AOB by Fe^3+^ flocs likely explains the spatial delay of NH ^+^ removal in full-scale sand filters

Fe^3+^ flocs are the main inhibitor of NH_4_ ^+^ oxidation in sand filters. We observed this phenomenon at laboratory-(Figure 5) and full-scale (Figure 7), and our results support the pioneering work of de Vet et al., (2009), which concluded that microbial nitrification in trickling filters might decline due to the irreversible accumulation of iron deposits. Likewise, the detailed molecular study of Tong et al., (2022) reported that Fe^3+^ flocs formed via homogeneous Fe^2+^ oxidation attach to the cell surface of *Pseudomonas putida*, severely decreasing its Mn^2+^ removal capacity. Nitrification kinetics in our lab-scale sand filter columns were fitted to a first-order rate equation and gave NH ^+^ oxidation rate constants (54-80 h^-1^) slightly above those found by Lopato et al (2013) (8-16 h^-1^) and references therein (9-17 h^-1^) in full-scale systems, likely due to the loss of efficiency ascribed to flow pattern heterogeneity at bigger scales (33). After 4 days of Fe^2+^ dosing, NH ^+^ removal capacity completely stopped at the filter top (NH_4_ ^+^ removal rate constant ∼ 0) (Figure 5). Our experiments do not allow us to rule out completely the contribution of mass transfer limitations to the spatial delay in NH_4_ ^+^ removal in full-scale sand filters as a consequence of filter clogging but suggest that they do not play a major role (Table 1). On the contrary, Fe^3+^ floc contents in the batch test experiments were 1 to 2 orders of magnitude below the ones in lab-scale (Figure 4E) and full-scale filters (Figure S3), but were still high enough to decrease the maximum NH_4_ ^+^ removal capacity of AOB on sand grains by half (Figure 6). Taken together, these results provide solid evidence that (i) Fe^3+^ flocs inhibit the NH ^+^ oxidizing capacity of AOB and that (ii) this mechanism is likely the main responsible for the spatial delay of NH_4_ ^+^ removal in full-scale sand filters. The exact physiological mechanisms remain to be elucidated.

**Figure 5:**
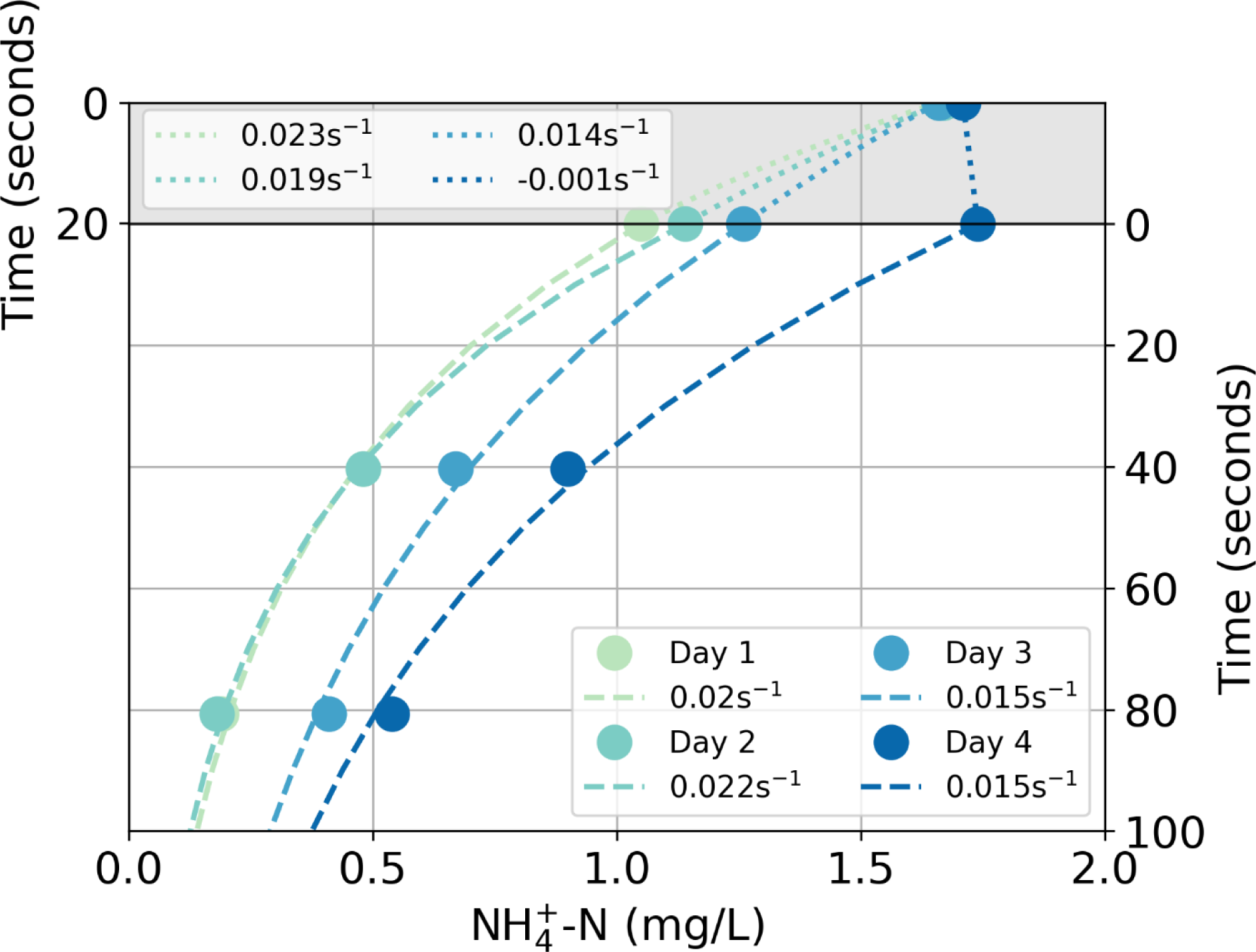
NH_4_ oxidation rate constants (k) estimated for the top 5 cm of the filter bed (gray background) and the downstream sections (white background. Dots show the observed concentration, and dashed lines show the fits used to estimate k values. R^2^ > 0.996 for all datasets.

**Figure 6.**
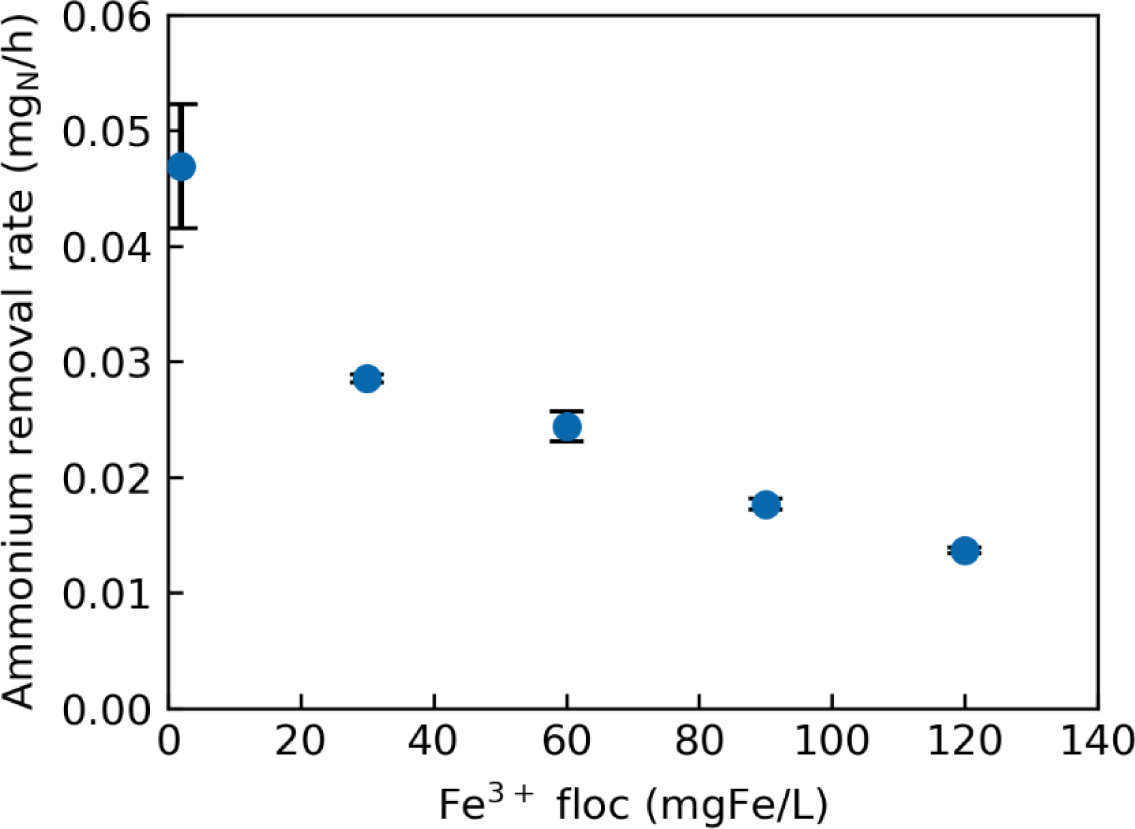
Maximum NH ^+^ removal rates of the sand grains with increasing amounts of Fe^3+^ flocs in the medium. Fe^3+^ flocs were added every 30 min during a period of 3h, mimicking the conditions of full-scale sand filter supernatants. X-axis indicates the concentration of Fe^3+^ at the end of the experiment. Activities were quantified in triplicates, error bars represent standard deviation between them. All data points included in Figure S4.

**Figure 7.**
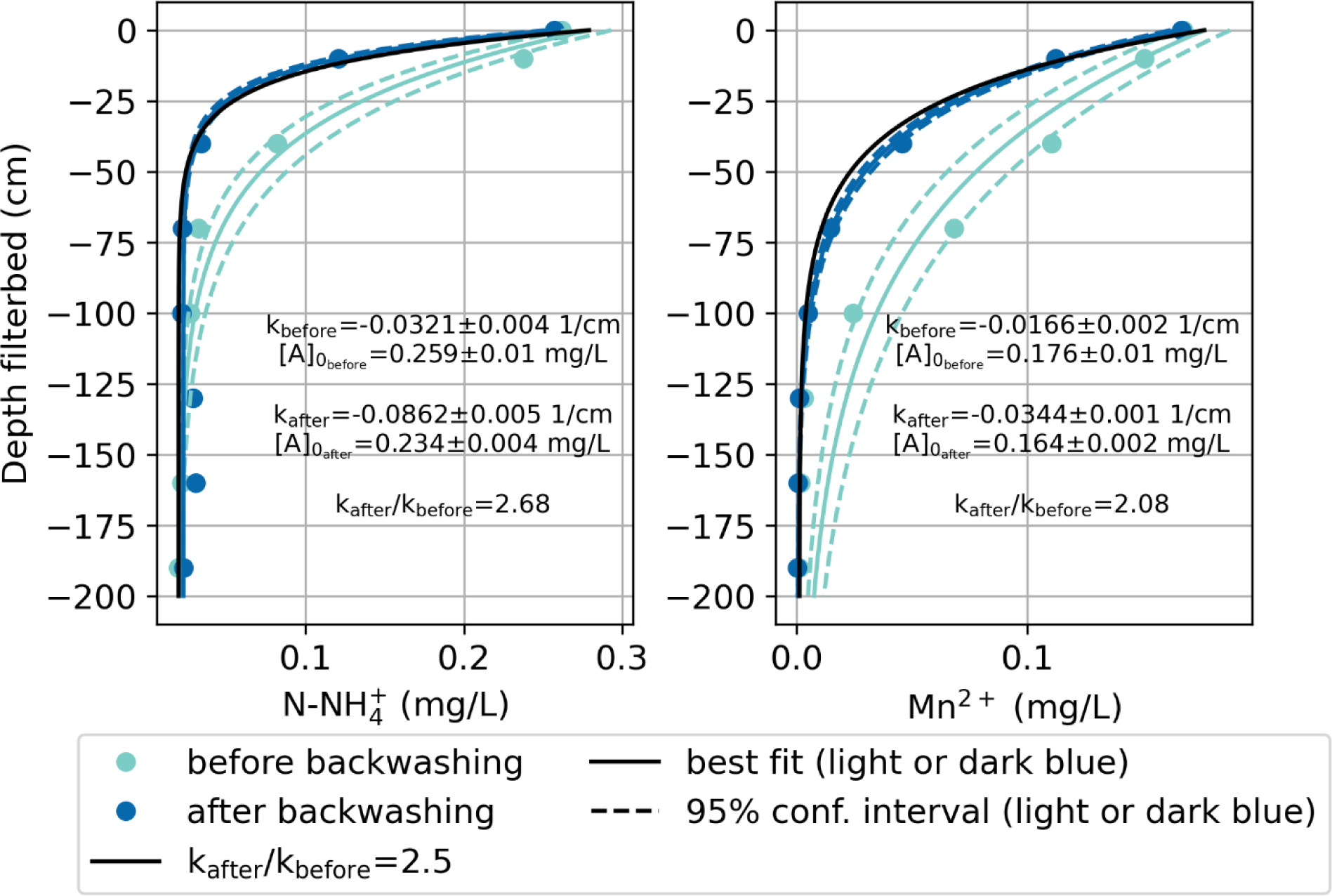
NH_4_^+^ and Mn^2+^ concentration profiles along the depth of a full-scale sand filter before (green) and after (red) backwashing. Lines indicate the fitting of the curves to first-order removal processes (full), and the 95% confidence interval of the fitting (dashed).

### 4.3. Ecological and industrial implications

Ammonium diffusion into the biofilm - *i.e.* mass transfer - has traditionally been considered to limit the nitrification capacity of sand filters fed with iron-free water (34,35). Yet, the latest evidence indicates that NH_4_ ^+^ removal is limited by biokinetics, the product of AOB abundance and its maximum specific NH_4_^+^ removal rate (36). In this work, we show that the presence of Fe^3+^ flocs inhibits the maximum NH ^+^ removal rate of AOB. Combined, these findings suggest that Fe^3+^ flocs effectively reduce the nitrification capacity of sand filters, and set avoiding Fe^3+^ floc formation as a clear optimization target. Strategies to segregate the removal of Fe and NH_4_ ^+^ have already been proposed and tested, and often encompass controlling the oxidation-reduction potential to avoid homogeneous Fe oxidation. Recent successful examples are anaerobic Fe^2+^ precipitation as vivianite (37), anoxic biological nitrate and Fe^2+^ co-removal (38) and aerobic biological Fe^2+^ oxidation at low oxygen concentration (39). Nitrification problems caused by Fe^3+^ flocs are currently bypassed because sand filters are usually over dimensioned (40). Nonetheless, finding solutions to directly tackle or avoid Fe^3+^ floc inhibition of nitrification is fundamental for the transition move towards resource-efficient, environmentally-friendly groundwater treatment. Beyond drinking water treatment, this work sheds light on the interplay between Fe and NH_4_ ^+^, two widespread and central compounds in many ecosystems across the planet.

## 5. Conclusions

In this work, we systematically and quantitatively evaluated the individual impact of the main Fe^2+^ oxidizing mechanisms and the resulting products on biological NH_4_ ^+^ oxidation in groundwater filters. Using a combination of batch essays, lab-scale columns experiments, full-scale column characterization and modelling, we conclude that:

- Dissolved Fe^2+^ and reactive oxygen species do not affect NH_4_ ^+^ oxidation
- Fe oxide coating aids sand grain colonization by NH_4_ ^+^-oxidizing bacteria
- Fe^3+^ flocs inhibit NH_4_ ^+^ oxidation by reducing the nitrifying capacity of AOB
- Changes in transport patterns due to clogging do not play a major role in NH_4_ ^+^ oxidation
- The inhibition of NH_4_ ^+^ oxidation is reversible and reduced by backwashing

Overall, our work sets finding solutions to directly tackle or avoid Fe^3+^ floc inhibition of nitrification as a priority to optimize groundwater treatment and transition towards the next generation of sustainable and efficient sand filters.

## 6. Authors contributions

FCR, EK, MLa, MvL, and DvH conceived the study. FCR, EK, SM, LV, and MLe conducted the experiments. EK developed the model. FCR, EK, SM, MLa, MvL, and DvH performed data analysis and/or helped interpret the results. FCR wrote the manuscript, with contributions from EK, MLa, MvL and DvH. All co-authors critically reviewed the manuscript and approved the final version.

## 7. Acknowledgements

The authors would like to thank Frank Schoonenberg (Vitens N.V) and Weren de Vet (WML Limburgs Drinkwater) for the full-scale data and thorough discussions This work was financed by the NWO partnership program ‘Dunea–Vitens: Sand Filtration’ (project 17830) of the Dutch Research Council (NWO) and the drinking water companies Vitens NV and Dunea Duin & Water. ML (VI.Veni.192.252) and EK (project 18369) were supported by NWO.

## 8. Supplementary information

### 8.1. Concentration profile DWTP Hammerflier

**Figure S1.**
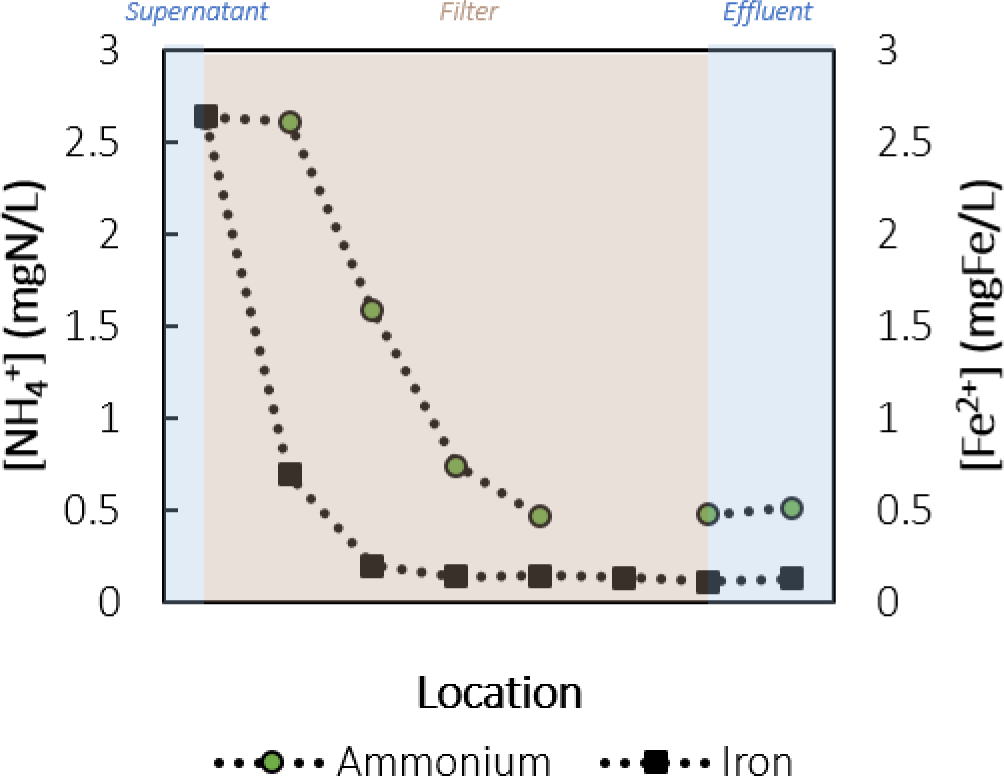
NH_4_^+^ and Fe^2+^ concentration along the height of a full-scale filter in DWTP Hammerflier. NH_4_^+^ removal is absent at the filter top and only starts when Fe^2+^ is oxidized. Filter media samples from the top were used for the column experiments.

### 8.2. Clogging affects hydraulic conditions during column experiments

Hydraulic conditions changed during the lab-scale sand filter experiments (Figure 4 and 5), as a result of clogging with Fe^3+^ flocs. Supernatant water levels increased over time, which showed that the clogging increased the resistance in the column. These hydraulic changes were also observed in the breakthrough curves of the salt spike tracer tests (Figure S2A), which were performed daily. The breakthrough curves showed more tailing over time and represented a wider variety of residence times of the influent water. Furthermore, the salt spike seemed to arrive a little earlier after clogging, but, note, that this deviation was always <1 minute (S2B).

**Figure S2:**
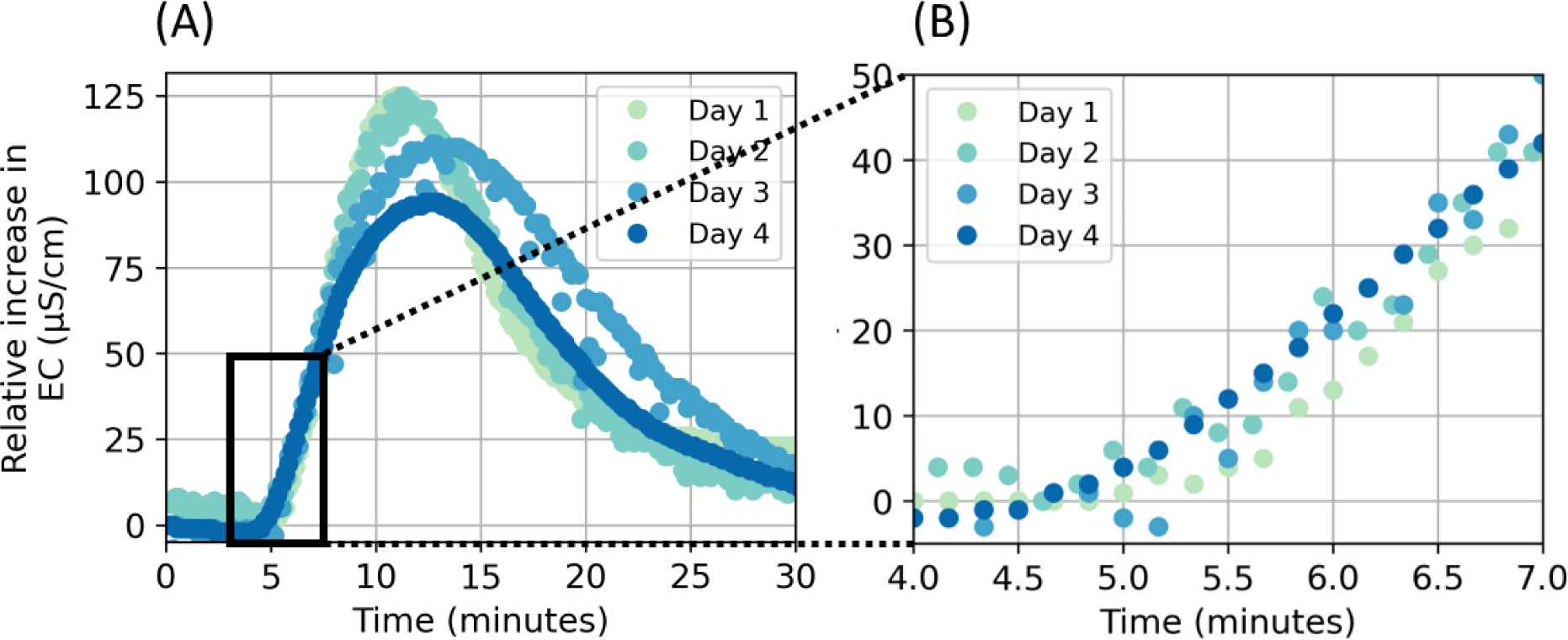
The relative increase of conductivity measured in the column during the daily performed salt spike tracer tests, wherein (A) the full breakthrough is shown and in (B) the arrival of the salt spike is highlighted.

### 8.3. Dissolved and particulate Fe before and after backwashing in full-scale filter

**Figure S3.**
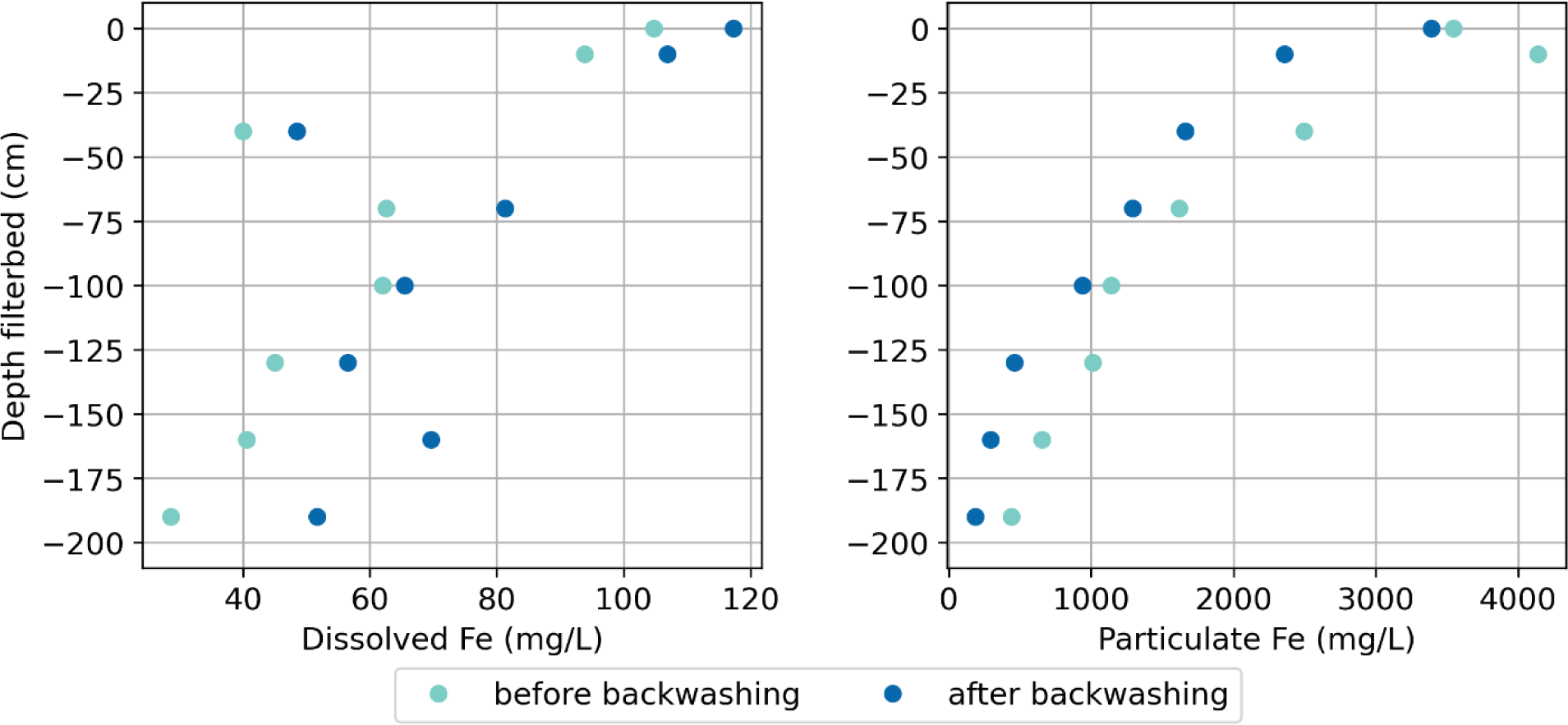
Dissolved (Fe^2+^) (A) and particulate (Fe^3+^ flocs) (B) Fe concentration before (green) and after (red) backwashing along the depth of a full-scale sand filter in drinking water treatment plant Hammerflier.

### 8.4. Batch test data for Fe^3+^ effect on NH_4_ ^+^ removal rate

**Figure S4.**
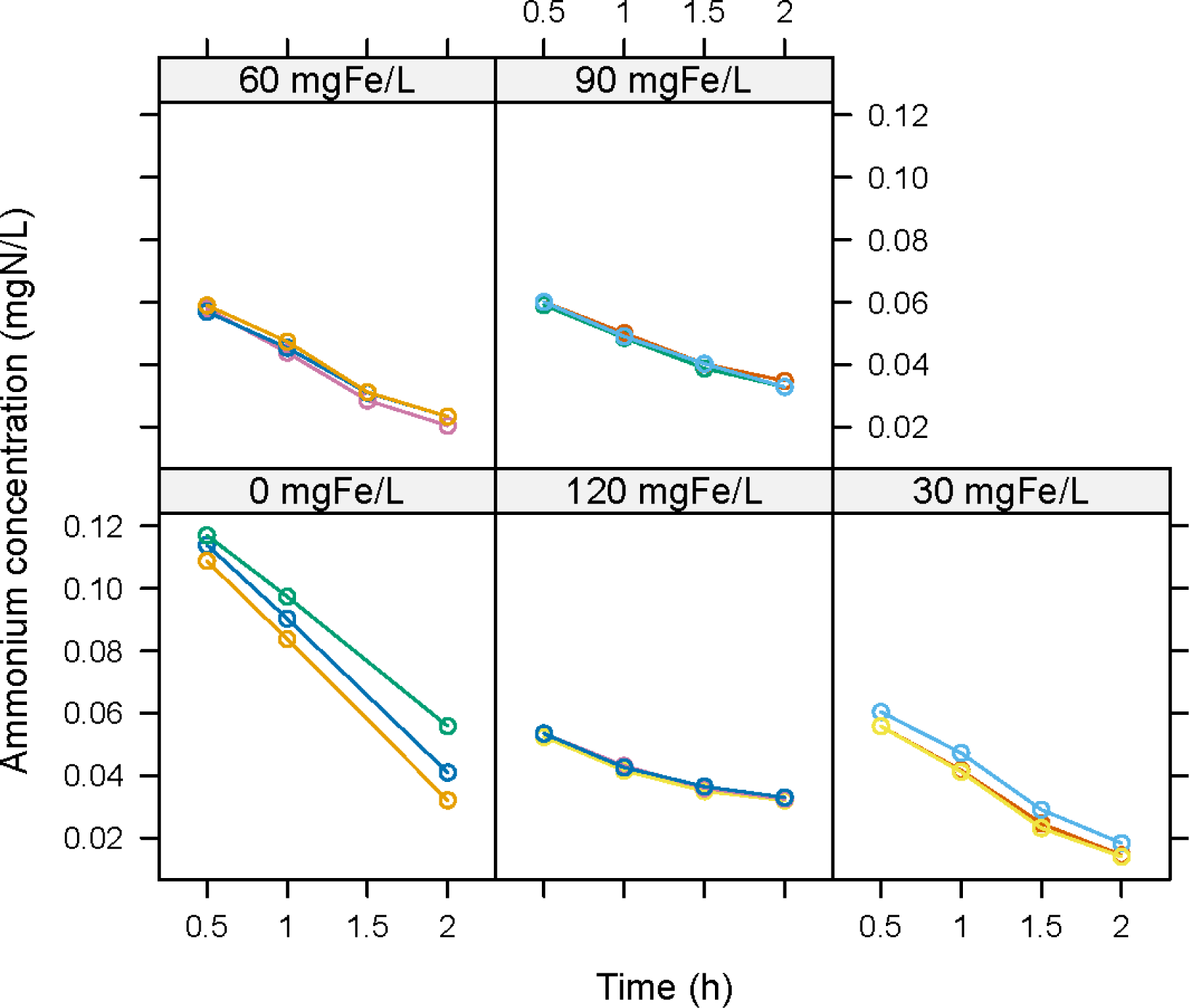
NH_4_^+^ concentration vs time in batch tests at 0, 30, 60, 90, and 120 mgFe flocs/L. Each line represents a different batch test.

